# Efficient strategies for screening large-scale genetic interaction networks

**DOI:** 10.1101/159632

**Authors:** Raamesh Deshpande, Justin Nelson, Scott W. Simpkins, Michael Costanzo, Jeff S. Piotrowski, Sheena C. Li, Charlie Boone, Chad L. Myers

## Abstract

Large-scale genetic interaction screening is a powerful approach for unbiased characterization of gene function and understanding systems-level cellular organization. While genome-wide screens are desirable as they provide the most comprehensive interaction profiles, they are resource and time-intensive and sometimes infeasible, depending on the species and experimental platform. For these scenarios, optimal methods for more efficient screening while still producing the maximal amount of information from the resulting profiles are of interest.

To address this problem, we developed an optimal algorithm, called COMPRESS-GI, which selects a small but informative set of genes that captures most of the functional information contained within genome-wide genetic interaction profiles. The utility of this algorithm is demonstrated through an application of the approach to define a diagnostic mutant set for large-scale chemical genetic screens, where more than 13,000 compound screens were achieved through the increased throughput enabled by the approach. COMPRESS-GI can be broadly applied for directing genetic interaction screens in other contexts, including in species with little or no prior genetic-interaction data.

## Introduction

Systematic mapping and analysis of genetic interaction networks is a powerful means of characterizing gene function and provides a valuable resource for exploring the relationship between genotype and phenotype. A genetic interaction occurs when mutations combine to generate an unexpected phenotype [1]. A negative interaction (e.g. synthetic lethality) results when different mutations, neither lethal individually, combine to cause cell death [2]. Conversely, positive interactions occur when mutations produce a phenotype that is less severe than expected [1]. Several organisms have been systematically screened for genetic interactions, including the budding yeast, *Saccharomyces cerevisiae*, where analysis of almost all possible 18 million double mutant combinations led to an extensive genetic interaction network consisting of ∼550,000 negative and ∼350,000 positive interactions and representing over 90% of all yeast genes [3]. This near-complete network in S. cerevisiae provides a powerful basis for understanding the basic principles of genetic interactions and developing more efficient strategies for screening them in other contexts.

Genome-wide analyses in yeast highlighted key properties of genetic interactions. For example, genes that function as part of the same biological pathway or protein complex tend to share similar patterns of negative and positive genetic interactions [3, 4]. Thus, the set of genetic interactions for a given gene, termed a genetic interaction profile, provides a rich and quantitative phenotypic signature indicative of gene function [3, 4]. These quantitative genetic interaction profiles can be assembled into a network in which genes with similar interaction patterns are located next to one another while genes sharing less similar interaction profiles are further apart in the network. The resulting global network based on genetic interaction profile similarity revealed a hierarchy of modules reflecting the functional architecture of a cell and provided a powerful resource for predicting gene function [3].

The global map of digenic interactions amongst loss-of-function mutations in haploid yeast is but one representation of the complex multi-dimensional genetic landscape. It is also important to understand the general principles associated with genetic interactions involving gain-of-function alleles and more complex genetic interactions that can occur in diploid and polyploid organisms [5–7], across a variety of different cell types of metazoans and within whole animals [8–10], or between hosts and their symbiotic organisms [11] [12]. Furthermore, while it is clear that genetic interactions can be conserved from yeast to humans [13, 14], we still lack an understanding of how global networks evolve and how a specific network is modulated in response to environmental or genetic background effects [15, 16]. Importantly, genome-scale application of CRISPR-Cas9 genome editing approaches offer the potential to map analogous genetic interaction networks in human cells [17–19]. However, mapping near complete genetic networks, analogous to the one constructed for *S. cerevisiae*, under diverse temporal and environmental conditions is resource intensive and often technically infeasible, particularly for higher eukaryotic cells and organisms with more complex genomes. Thus, development of rational and scalable strategies is required to optimize screening that can be generally applied to efficiently map genetic interaction networks for different organisms under a variety of experimental contexts.

While optimization strategies have been reported for mapping protein-protein interactions, these methods do not readily apply to genetic interaction networks [20]. The unique properties of genetic interaction networks warrant a systematic study of different approaches and the development of new methods. Previous studies identified several functional, physiological and evolutionary properties associated with genetic interaction frequency in yeast [3, 4, 21]. For example, perturbation of genes that result in increasingly severe fitness defects participate in more negative and positive interactions in the global yeast genetic network [3, 4]. Indeed, a model based on gene-specific properties derived from the *S. cerevisiae* genetic network successfully identified highly connected genes (i.e. genetic interaction network hubs) in a distantly related yeast species [21]. Another method prioritized genes based on minimum uncertainty in cluster membership based on a clustering of genetic interaction data [22]. Although these approaches can identify genes that display many genetic interactions, it is not clear if the genetic interaction screens based on subsets of genes identified by these methods are able to capture the structure, topology and functional spectrum of a complete, genome-wide genetic interaction network

In this paper, we present novel and generalizable methods for identifying prioritized subsets of genes with genetic interaction profiles that can recapitulate a global genetic interaction network. To demonstrate the impact of screen prioritization, we apply one of our approaches to the problem of optimal selection of screens for functional profiling to support a large-scale chemical genetic screen. Optimization in this context enabled us to achieve a nearly 10-fold speed-up in the rate at which chemicals were profiled against a optimally selected mutant collection while retaining, and even improving, the amount of information extracted from the screen.

## Results

### Defining an objective for optimizing genetic interaction screens

There are several possible objectives one might have in screening genetic interactions, and the optimal strategy depends on the goal of the screen. Perhaps the simplest objective is to discover new individual genetic interactions at the fastest rate. More specifically, in the context of digenic genetic interactions, one objective could be to select pairs for screening in each round that maximize the number of previously undiscovered genetic interactions. For this simple objective, an understanding of the hubs in the network, i.e. the highly-connected nodes, provides an efficient strategy for this objective. For example, in the two different yeast species in which the largest GI screens have been completed, *S. cerevisiae* and *S. pombe*, it has been reported that one of the primary correlates of the number of genetic interactions for a given gene is the extent of the phenotype (for example: fitness defect) associated with the individual allele [4, 21]. In fitness-based screens, single mutants with stronger fitness defects tended to be strongly associated with larger number of genetic interactions (r = 0.73), even after accounting for the expected multiplicative effect of the single mutant [4]. This strong correlation with the strength of the single mutant phenotype suggests the simple but effective strategy of selecting mutants for screening in order of the severity of their single mutant phenotypes. We measured the effectiveness of this approach on the comprehensive version of the *S. cerevisiae* genetic interaction network, consisting of ∼5 million screened pairs [4]. Indeed, prioritizing mutants for screening based on the strength of the single mutant phenotype produces new genetic interactions at a rate substantially faster than a baseline approach of random selection of genes for screening (S1a Fig), and only slightly below the maximum possible rate of detection (S1a Fig). For example, screening 25% of the genome based on low-fitness uncovers 60% of the interactions (S1a Fig). This strategy could be refined even further if one leverages more complex predictive models for the degree of genetic interactions associated with a given gene. For example, Koch *et al.* report a collection of physiological and evolutionary properties, in addition to the strength of the single mutant phenotype that are predictive of genetic interactions, and notably, this model appears to work in multiple distantly related species [21]. This simple strategy of predicting hubs based on readily accessible features provides a reasonable basis for optimizing screens when the goal is simply to recover new interactions as quickly as possible.

A more specific objective than simply detecting new genetic interactions is generating genetic interaction profiles that enable the characterization of gene function. Several previous studies have demonstrated that a primary strength of genetic interactions is that a gene’s profile, i.e. its signature of interactions across a collection of mutants, can be compared with other genes’ profiles to identify genes that function in the same specific bioprocess, pathway, or even protein complex. Thus, an important objective in screening genetic interactions is to select a set of mutants for screening that will produce an interaction profile that most efficiently characterizes gene function. The remainder of our study is focused on optimizing this objective.

Before designing a method to tackle this problem, we first evaluated the potential feasibility of compressing genetic interaction profiles. More specifically, we measured the extent to which screening genes against an increasing, randomly selected, subset of the genome could capture the information contained in a complete genetic interaction profile. The functional information captured by a genetic interaction profile was measured by the ability genetic interaction profile similarity (i.e. pairs of genes exhibiting highly similar genetic interaction signatures) to predict functionally related gene pairs as measured by co-annotation to Gene Ontology (GO) terms. Encouragingly, we observed that the performance of genetic interaction profiles in predicting functionally related genes increased dramatically with profile length and that even a randomly selected subset of genes of could achieve relatively strong performance (S1b Fig). For example, in *S. cerevisiae* screening 500–1000 genes produces near saturation level of performance (S1b Fig). This dramatic increase in performance with only a small, randomly selected subset of the genome is likely the result of the fact that genetic interactions are highly structured, connecting between and within functional modules rather than at the level of individual genes [3, 4, 23], and thus there is a high degree of redundancy in the information captured by any single screen. This conclusion that genetic interaction profiles can be significantly compressed is also supported by a standard PCA analysis, which confirms ∼80% of the variance in the GI network can be explained by 500 principal components (S2 Fig). Given the encouraging performance of a randomly selected, small subset of genes in capturing functional information in GI profiles, we reasoned that an intelligent approach to screen selection could further improve the amount functional information captured in screened profiles. We discuss algorithms that specifically address this problem in the sections that follow.

### COMPRESS-GI: A novel gene selection method for optimizing genetic interaction profile similarity

Towards the goal of selecting a small set of informative genes optimized for measuring functional relationships based on genetic interaction profile similarity, we developed a novel algorithm, COMPRESS-GI (**COM**press **P**rofiles **R**elated to **E**pistasis by **S**electing **I**nformative **G**enes). We developed two different versions of algorithm, batch (COMPRESS-GI) and iterative (iCOMRESS-GI), to address two different scenarios. The batch version is designed for instances in which one already has access to a large collection of genome-wide genetic interaction screens and wishes to use this collection to select a small set of mutants that captures most of the information contained within the already collected profiles (e.g. to perform increased throughput screens with a reduced set). The iterative version is designed for the scenario in which few or no genetic interaction screens have been completed and provides a iterative strategy for query selection.

### COMPRESS-GI

The batch version of COMPRESS-GI leverages existing genetic interaction data consisting of genome-wide interaction profiles for at least a few hundred genes across the genome and known gene relations from the Gene Ontology (GO) to define an informative set of diagnostic genes that is informative for discovering functional relationships. Given several hundred genome-wide genetic interaction profiles, the goal of this algorithm is to recapitulate genetic interaction profile similarity using a minimal number of mutants selected from the genome-wide profiles. More specifically, COMPRESS-GI will select a subset of the genes such that the similarity network generated using only profile derived from interactions with the selected genes can be used to effectively predict functionally related pairs. The performance of the GI profile similarity network derived from a subset of genes can be quantified using precision-recall characteristics based on a standard of co-annotation to the Gene Ontology (Fig 2a; see Methods) [24]. Based on this objective, we use a step-wise exhaustive greedy approach, where we select the most informative gene as measured by precision-recall performance of the similarity network generated based on profiles defined by interactions with each single gene. We continue iteratively maximizing the precision-recall performance of the resulting profile similarity networks by adding genes that maximally increase the performance when added to the already selected set (Step 2, Fig 2b; see Method for details). However, this process is potentially biased by the starting gene, so we repeat the process with several different starting genes, each taken from the top 50 most informative based on the single gene’s profile (Step 1). In addition, we know from previous experience that precision-recall performance measured based on a global GO standard can be dominated by genes belonging to one or few functional categories [24]. Thus, we repeated the process described above independently for each of 13 major functional categories, each time focusing the GO co-annotation standard to retain positive co-annotations belonging only to the broad functional category of interest while removing other positive co-annotations (Step 3; see Methods for details). Each one of these runs, focused on a different functional neighborhood of interest, resulted in a ranked list of genes that optimized recovery of functionally related gene pairs based on compressed GI profiles. We combined these lists of informative sets of genes to obtain a single, global list of around 200 genes (Step 4).

**Fig 1.**
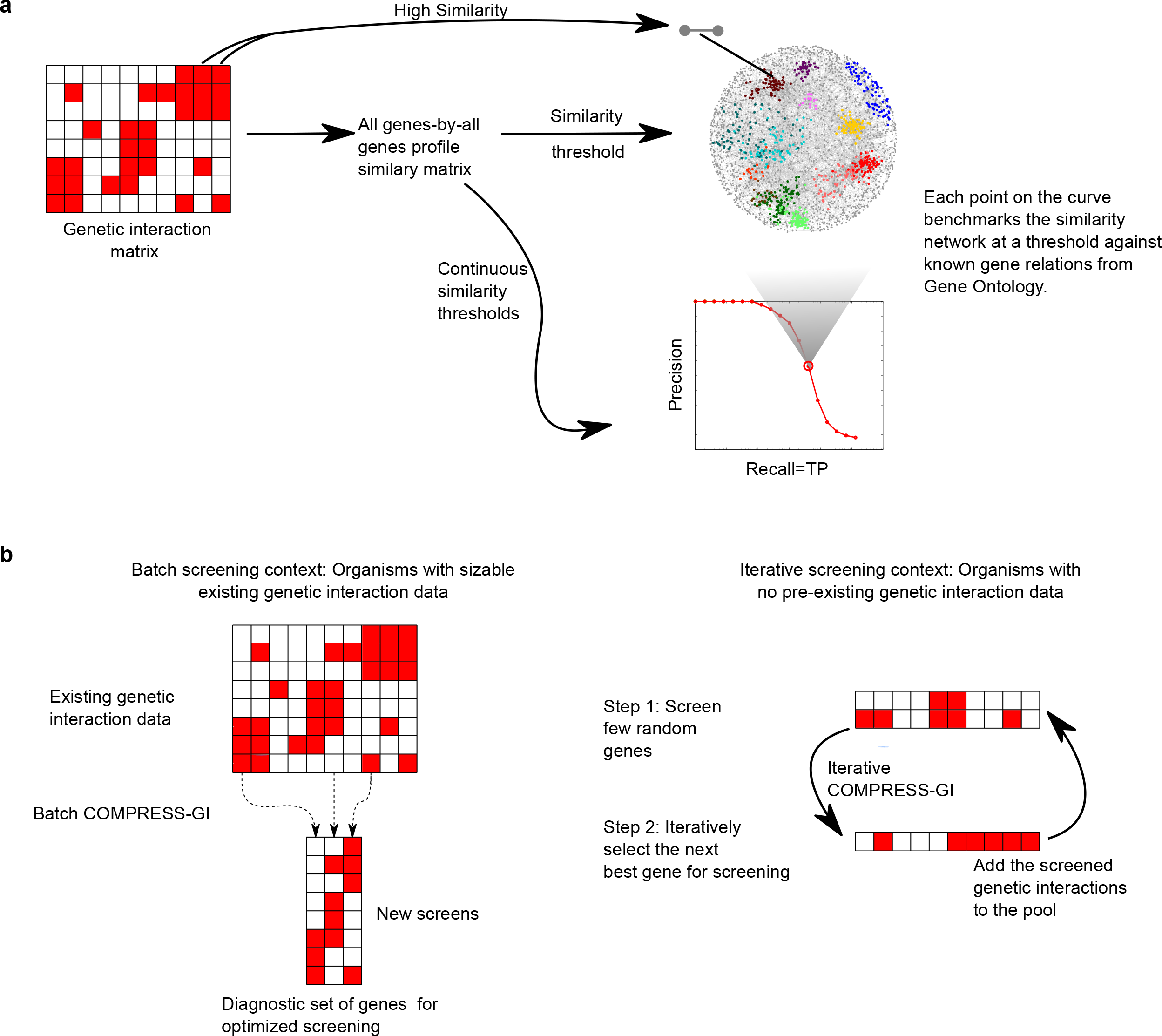
Overview of genetic interaction profile similarity objective. (a) The COMPRESS-GI algorithm maximizes the information content in the genetic interaction profile similarity network using precision-recall metric. Given a profile similarity network at a defined similarity threshold and using the similarities as predictions, the similarities can be compared with positive and negative co-annotation based on an external standard (e.g. the Gene Ontology) to measure precision and recall. We address two different screening scenarios with the COMPRESS-GI algorithm: (b) The batch version of COMPRESS-GI is designed for scenarios in which substantial genetic interaction data already exist while the iterative version of COMPRESS-GI is designed for screening genetic interactions in contexts with little or no pre-existing genetic interaction data.

**Fig 1.**
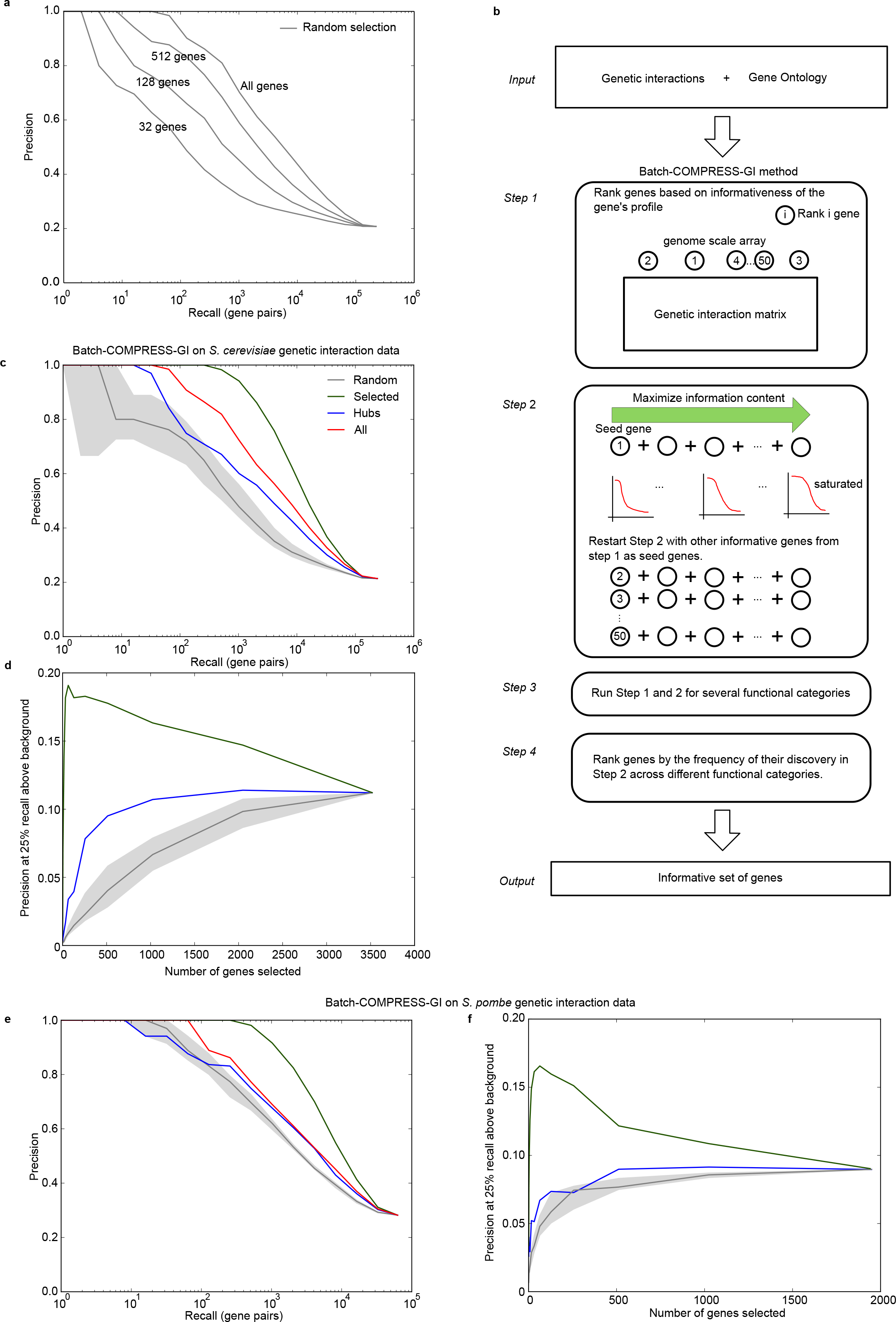
COMPRESS-GI algorithm and evaluation. (a) Precision-recall curves reflecting the predictive power of the genetic interaction profile similarity network generated with different random subsets of the profile. The precision-recall curves shown are the mean of 11 different random runs for each gene set size mentioned. (b) A flowchart describing the COMPRESS-GI method. The inputs for the COMPRESS-GI algorithm are genetic interaction data, a Gene Ontology or other standard for gene pair functional relationships, and gene membership in broad functional categories. The algorithm outputs an informative set of genes, which is a small subset of the original screened set against which profiles are maximally informative. (c) Precision-recall evaluation of the top 100 COMPRESS-GI selected genes and comparison with an equal number of randomly selected genes, equal number of top hubs, and the entire S. cerevisiae genetic interaction dataset. (c) The evaluation in (b) was repeated for different functional categories by modifying the Gene Ontology standard, and precision at 25% recall was averaged across the different functions. The randomization for (c) and (d) was repeated 100 times. The middle line is the median across those runs, and the boundaries of the shaded area represent the first and third quartile, respectively. (e,f) shows analyses similar to (c,d) conducted on *S. pombe data*.

**Fig 2.**
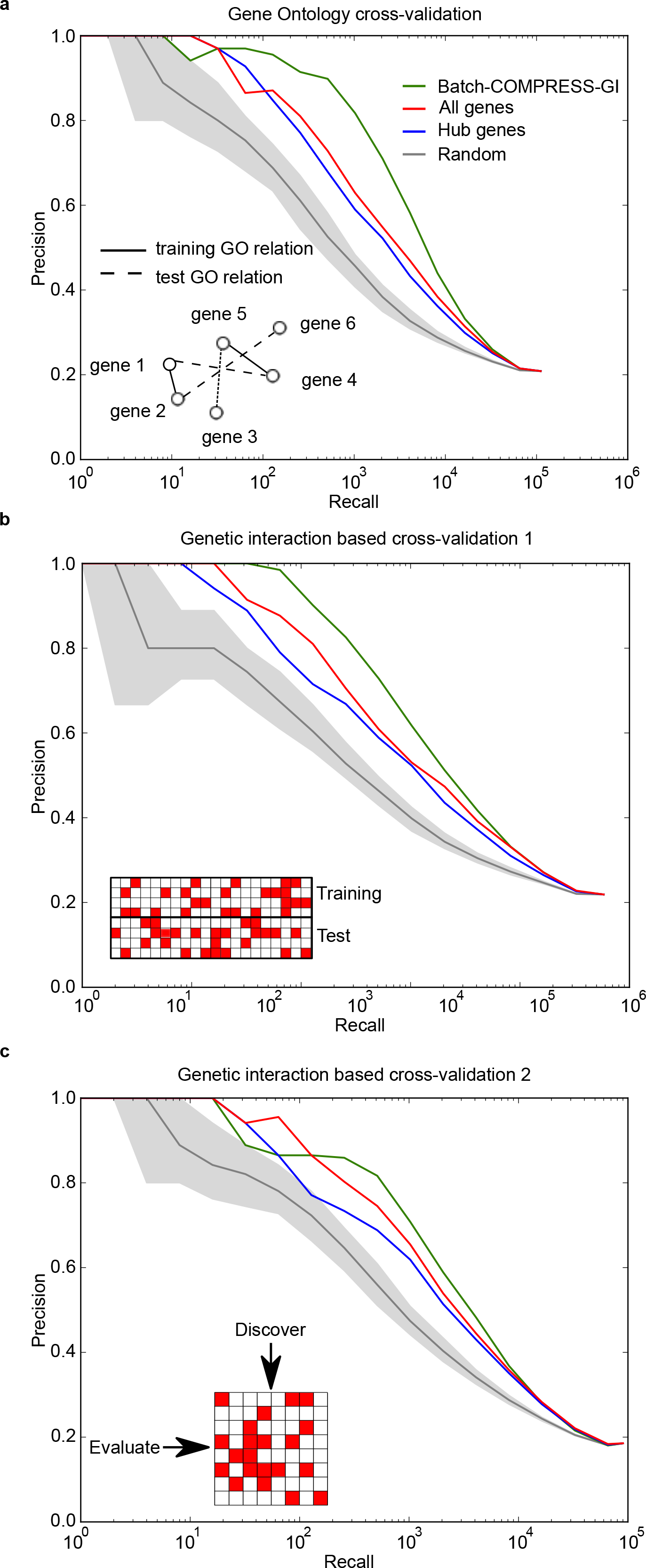
Cross-validation of the COMPRESS-GI algorithm. (a) The Gene Ontology is split into training and test samples for cross-validation, and COMPRESS-GI is run on the training GO standard, and the informative set of genes is discovered. The informative set of genes is evaluated using precision-recall measures on the held-out part of the GO standard. (b) Similarly, the cross-validation is repeated on different parts of genetic interactions data by splitting it into training and test halves on the query side. (c) The informative set of genes discovered on the training genetic interaction data is evaluated on the test half. The utility of the informative set of genes discovered on the array side is tested on the query side. The square genetic interaction matrix, which has the same genes on both the array and the query sides, is used for cross-validation in (c). The hubs in each of the plot are dataset-specific. For example, in (b), hubs from the test genetic interactions are used. For the random baselines, the randomization was repeated 100 times, and median precision-recall performance is plotted. The lower and upper bounds of the gray area represent the first and third quartile across these randomizations.

### Evaluation of the COMPRESS-GI algorithm

The set of genes selected by the COMPRESS-GI algorithm provides a substantial improvement in performance as compared to a random selection of screens. The precision-recall characteristics of the top 100 selected genes is significantly better than even the 75th percentile from several random runs where an equivalent number of mutants was selected randomly (Fig 2c). To further understand the performance of the selected set, we evaluated the COMPRESS-GI genes for ability to predict gene-relations within specific functional categories. When we compare the aggregated precision at 25% recall across 13 functional categories and across different number of selected genes, we see that the COMPRESS-GI selected genes consistently perform better than the random baseline (Fig 2d). The performance improvement over randomly selected genes is even more striking for smaller sets of genes, suggesting that our method of intelligent selection of informative genes with little redundancy cannot be easily achieved through random sampling. Beyond the random selection baseline, we also compared our approach to a reasonable strategy of selecting high-degree (hub) genes and observed that our selected set of genes also performed better than an equal number of hubs (Fig 2c,d). More interestingly, by the chosen precision-recall metric, the selected genes perform better than the complete set of genes based on a global GO evaluation standard (Fig 2c), and comparably to the complete dataset on the function-specific evaluation (Fig 2d). We note that precision-recall performance does not monotonically increase with the number of genes selected. The better performance of the selected genes over the complete dataset likely suggests that there are non-informative or potentially low-signal genes in the complete dataset that detract from functional information in the genetic interaction data and thus, bring the precision-recall performance down relative to a smaller, intelligently chosen, set of informative genes.

Given our use of the GO functional standard during gene selection, to ensure that the COMPRESS-GI algorithm was not overfitting, we conducted two separate cross-validation experiments. First, we identified informative genes by applying COMPRESS-GI to only 50% of the existing genetic interaction screens (randomly selected) and then tested the ability of these selected genes to provide profile information for the held out 50% of the genetic interactions screens. In a second cross-validation experiment, we held out 50% of the GO annotations when selecting an informative gene set, and then benchmarked the selected set for its ability to recover held-out pairwise GO co-annotations. In both cross-validation experiments, the COMPRESS-GI selected genes were equally informative on held-out data and/or annotations (Fig 3a,b), suggesting that the approach is not overfitting.

**Fig 3.**
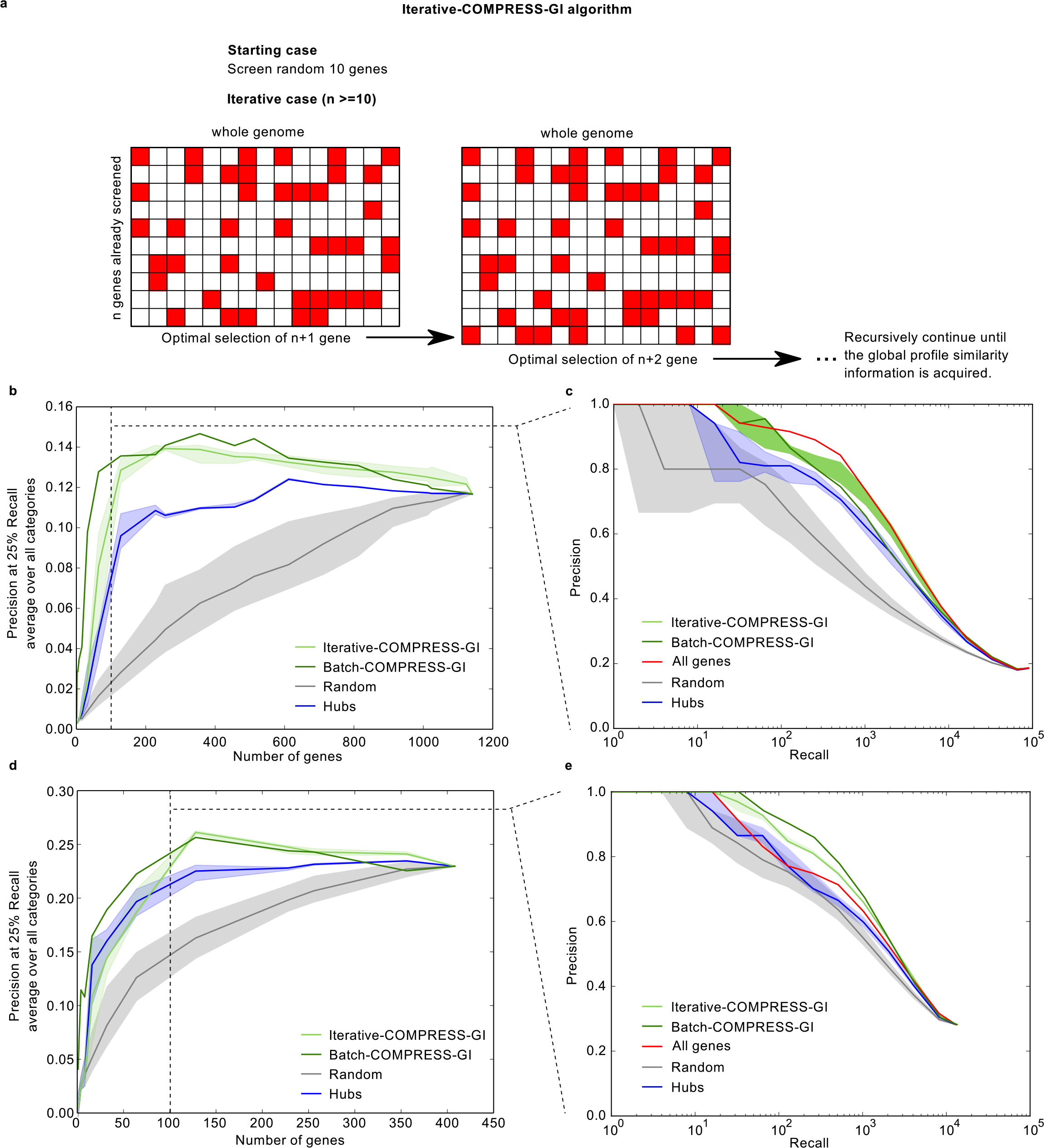
Iterative genetic interaction screening scenario. (a) The schematic describes the iterative genetic interaction screening scenario, which assumes no initial genetic interaction data other than 10 random screens for initialization. (b,c) The iterative COMPRESS-GI method is evaluated by simulating iterative genetic interaction screening on the square genetic interaction Costanzo *et al.* data. (b) compares the precision-recall curves for 100 genes selected by this approach with an iterative hub and random baseline approaches. (c) compares the precision at 25% recall performance of the different approaches averaged across all functional categories. In addition to these approaches, the performance of the complete genetic interaction data has also been added in (b,c). Since there is random variation due to the selection of the first 10 genes, the approaches were repeated 10 times each with different initial gene sets. Each of the random cases, where the rest of the 90 genes are random, was also repeated 10 times (d, e). Same evaluation as in (b,c) repeated on *S. pombe* genetic interaction data.

In a final evaluation, we explored the effect of the asymmetry of the genetic interaction screening platform on the performance of our algorithm. Synthetic Genetic Array (SGA) analysis, the technology that was used to produce the global genetic interaction map for yeast requires that in a given screen, a single query strain be crossed into an ordered array of second mutant strains [25, 26]. To verify that the performance of the genes we selected with COMPRESS-GI was not dependent on the configuration in which they were screened (e.g. as a query strain or an array strain), we limited our analysis to the subset of genes in the yeast genetic interaction network that appeared on both the query and array sides of the interaction matrix [4]. We then discovered informative sets of genes by running the COMPRESS-GI on the array side checked the information content of the same set genes on the query side, and vice versa. Indeed, we confirmed that the selected genes were informative in both cases, suggesting that our approach is able to select truly informative genes and is not sensitive to the assymetry of the screening platform (Fig 3c).

### iCOMPRESS-GI: an iterative approach for screening genetic interactions

The iCOMPRESS-GI is a variation of the batch version where the method does not require a large set of pre-existing genetic interaction data to begin, and therefore, is more appropriate when limited or no genetic interaction data have been produced. The iCOMPRESS-GI was developed to address the limitation of the batch COMPRESS-GI method in that it requires a sizeable genetic interaction matrix as an input. The algorithm will be useful for prioritizing genes for genetic interaction screening in new organisms or for new conditions for species with already established genetic interaction networks. The iCOMPRESS-GI can be started with as few as 10 initial genome-wide genetic interaction screens, after which the algorithm will iteratively discover the next informative gene to be screened. This iterative process can be repeated after each additional gene is screened to obtain the next gene to be screened (Fig 4a).

The iCOMPRESS-GI approach approximates the batch COMPRESS-GI approach but is orders of magnitude faster (see Methods). Briefly, iCOMPRESS-GI optimizes the sum of products of similarities between genes and the known GO co-annotations between them (0, 1, or −1), which can be summarized as a Hadamard’s product or element-wise matrix multiplication. Using properties of the trace on a Hadamard’s product along with the cyclic property of the trace product of matrices, the problem can be reduced to a simple 0-1 knap-sack problem, giving each gene a score that is related to the gene’s informativeness (see Methods for derivation). The genes can be ranked by their scores and top genes can be selected and screened.

Given its computational efficiency, the iCOMPRESS-GI approach is suited for the iterative genetic interaction screening scenario where screens are selected in an online fashion after each additional screen. For comparison, we have also implemented a baseline approach, called “iterative hubs”, which is based on screening the highest-degree previously unselected hub after each screen. Both iCOMPRESS-GI and the iterative hub method are initialized based on genome-wide interaction screens of 10 randomly selected genes.

We simulated and evaluated the scenario of iterative genetic interaction screens on the Costanzo *et al.* 2010 genetic interaction data [4]. We selected a submatrix of the complete genetic interaction network such that genes on the array side were also on the query side, ensuring that we had screens for any genes selected from among the set of arrays, which resulted in a square matrix of 1141 query genes × 1141 array genes. As mentioned above, 10 genes were randomly screened first, followed by 90 iteratively selected genes, for a total of 100 query gene screens. To measure the performance of each approach, a profile similarity network was constructed by measuring similarity between all pairs of array genes based on the 100 selected query genes, and evaluated with the Gene Ontology co-annotation standard using precision-recall analysis. Similar simulations were conducted to select 100 genes using the baseline iterative hubs approach.

We observed that the iCOMPRESS-GI method performs better than both the iterative hubs approach and random screen selection in terms of the ability of the resulting profile similarity network to capture known functional relationships between genes (Fig 4b). We performed a side-by-side comparison with the batch COMPRESS-GI method with an equal number of genes, which serves as an upper bound for the iCOMPRESS-GI algorithm as the complete genetic interaction matrix is available at the outset of the batch algorithm. Gene selection was continued beyond 100 genes to the completion of the square matrix, and performance was evaluated across different functional contexts using precision-recall statistics as with the batch COMPRESS-GI. Again, the precision at 25% recall performance, averaged over the 13 functional contexts, is higher for the iterative COMPRESS-GI approach compared to iterative hubs and random (Fig 4c). The iCOMPRESS-GI performs exceptionally well relative to the baseline methods for selection of small sets of genes, making it especially useful for small studies in which less than ∼100 screens are feasible.

To test if our approach generalizes to species beyond *S. cerevisiae*, we carried out a similar simulation on the *S. pombe* genetic interaction network [27] (Fig 4d,e). As in our *S. cerevisiae* evaluation, we observed that the genes selected by the iCOMPRESS-GI approach perform better than both random and iterative hub baseline approaches. These positive results in both species suggest that the algorithm will be useful in other contexts as well.

### Application of COMPRESS-GI to enable large-scale chemical genetic screens

To demonstrate the utility of our COMPRESS-GI approach for optimizing genetic interaction screens, we applied it to facilitate a large chemical genetic screening effort. Chemical genetic screens involve screening a compound of interest against a collection of mutant strains with defined genetic perturbations [28–32]. Depending on the compound’s effect on a cell, individual mutants can be differentially sensitive and this profile of sensitivity measured across the mutant collection acts as a “fingerprint” that is indicative of a compound’s mode of action [32, 33]. This is a powerful unbiased approach that can be used to gain insight about the mechanism of action of uncharacterized compounds. Large-scale chemical genetic screening methods were pioneered in yeast [30–33] where much of the focus has been on screening chemicals against genome-wide mutant libraries.

Genetic interaction networks provide a key to deciphering chemical genetic profiles to identify specific molecular targets. Specifically, molecular targets of small molecules can be predicted using the idea that a compound’s behavior in a chemical genetic screen should mimic the behavior of a mutation in the corresponding target protein across the same mutant collection [33]. Chemical genetic profiles across a complete, genome-wide collection of mutant strains are ideal as they measure sensitivity or resistance with each gene mutant represented in the collection. However, if the major aim in collecting chemical genetic profiles is to measure profile correlation of candidate compounds’ profiles with genetic profiles or even profiles of other compounds, then chemical genetic screens against an intelligently selected subset of mutant should perform well at this task.

This is an ideal scenario for the batch version of the COMPRESS-GI algorithm. Given the near-complete genetic interaction network, our task is to select a small subset of single mutants against which candidate compounds should be screened to identify chemical-genetic interactions. These compressed genetic interaction profiles can then be compared to the complete database of genetic queries screened against the same subset of mutants to identify genes that have a similar profile of genetic interactions as the compound(s) of interest (Fig 5a). Given the fact that the interpretation of the chemical genetic interactions is being derived from profile similarities, our earlier analysis of the COMPRESS-GI algorithm suggests we could do this successfully with only a small fraction of the entire collection of mutants. For example, our application of the batch version of COMPRESS-GI resulted in the selection of ∼160 highly informative genes, comprising less than 5% of the non-essential deletion collection, but which recapitulated a profile similarity network capable of predicting known functional relationships as well as complete genetic interaction profiles. Importantly, reducing the number of mutants screened for chemical-genetic interactions could lead to significant resource savings. Fewer unique mutant strains require less growth media, and thus smaller volume of compounds is required, which is important when compound quantities are limiting. Furthermore, for next-generation sequencing based read-outs, as we use here [34], the throughput of the chemical genetic screening system is directly proportional to the number of unique mutants: reducing the number of mutants by 20-fold can result in a 20-fold increase in the number of compounds screened per lane of sequencing. Thus, conducting chemical genetic screens using a diagnostic mutant pool selected with COMPRESS-GI could substantially reduce the compound volume and overall cost.

**Fig 5.**
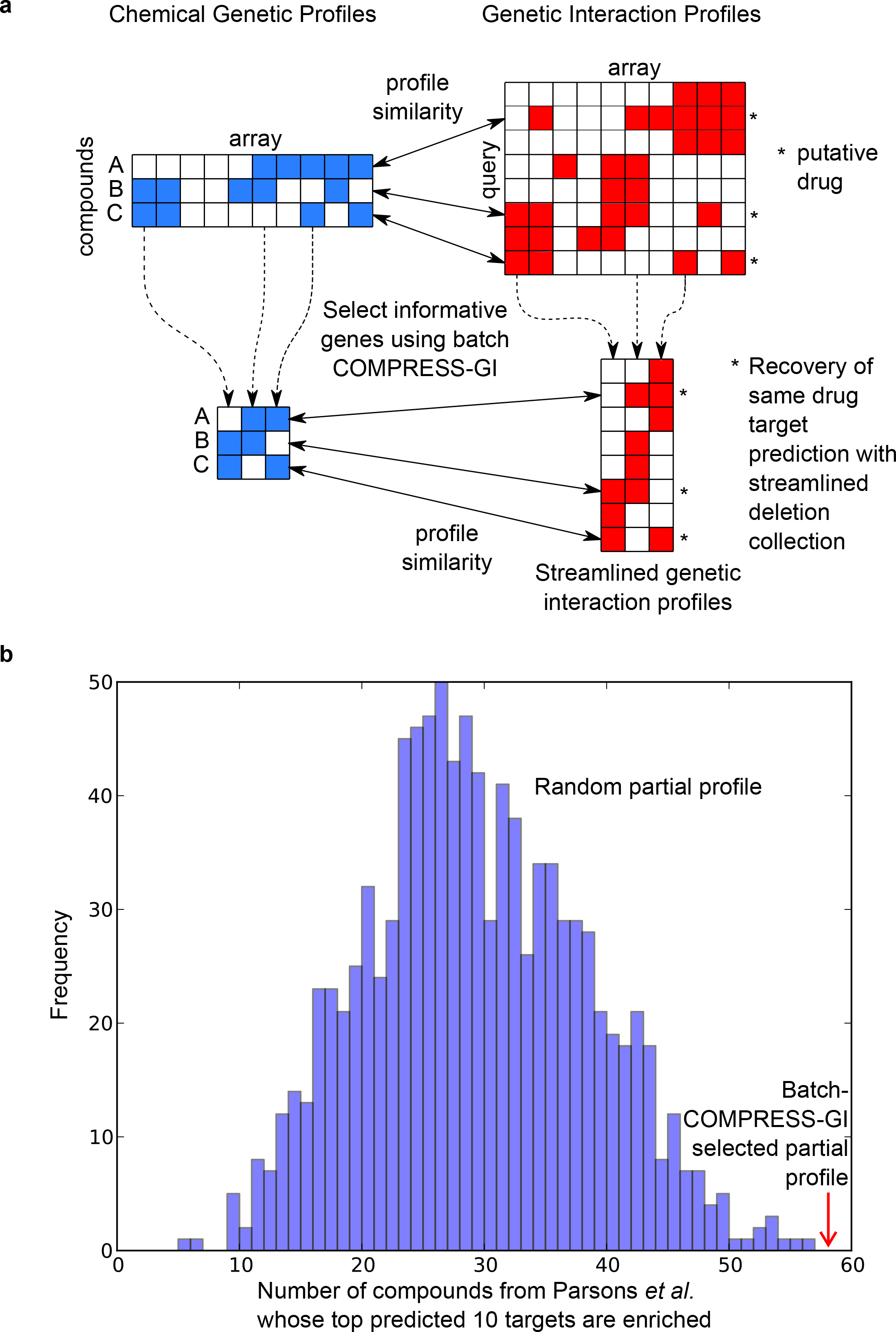
Application of diagnostic set of genes discovered by COMPRESS-GI to
chemical genetic screens. (a) The schematic illustrates that with COMPRESS-GI, diagnostic gene set selection, even with smaller partial profiles, we can recapitulate similar compound-gene target predictions. (b) The Fig compares the number of compounds out of 82 compounds in Parsons et al. study whose top 10 targets based on COMPRESS-GI selected genes (red arrow) with the random partial profile targets (blue histogram, random runs repeated 100 times) are enriched for Gene Ontology terms. The top targets of a compound are identified by finding query genes (rows) in the genetic interaction data that have maximum correlation with the chemical genetic interaction profile of the compound of interest.

#### In silico evaluation of compressed mutant pool for chemical genetic application

Before investing substantial experimental resources in chemical genetic screens using the diagnostic pool of mutant constructed with our COMPRESS-GI algorithm, we performed simulation studies with previously published chemical genetic interaction data to estimate the performance of the selected set of genes for chemical genetic applications. We leveraged data from a previous chemical-genetic study in which 82 compounds were screened for chemical genetic interactions against the entire collection of ∼4000 non-essential deletion mutants [33]. We compared the compound-target prediction capability of the compound profiles restricted to the 150 genes selected by COMPRESS-GI to the performance of compound profiles restricted to an equal number of randomly selected genes. As described earlier, compound-target prediction was conducted by finding genes with the most highly correlated genetic interaction profile to the chemical genetic profile for the compound of interest based on the assumption that the compound’s behavior will mimic the knock-out of the target gene [33]. We compared the enrichment of the top predicted targets using the diagnostic set of 157 strains with multiple runs of top predicted targets using a randomly selected set of strains of the same size. We found that the targets were more likely to be enriched for Gene Ontology (GO) terms when they were discovered from our COMPRESS-GI-selected set of mutants as compared to the same-sized randomly selected sets of genes (Fig 5b; p-value < 0.01). Significant enrichment of the top predicted gene targets indicates that the compound’s chemical genetic profile is consistent with a functionally coherent set of genes, which is most likely indicative of the bioprocessed affected by the compound. These results suggest that target predictions based on chemical genetic profiles against the diagnostic set of genes is more effective than equally sized sets of randomly chosen genes. Interestingly, chemical genetic profiles restricted to the diagnostic set of genes also performed better than the entire chemical genetic profile (59 vs. 42 of the 82 compounds showed GO process enrichment among the top predicted gene targets). These analyses suggest that the COMPRESS-GI-selected set of mutants indeed does perform well at capturing information in chemical-genetic profiles despite the fact that it was selected entirely based on genetic interaction screens.

#### COMPRESS-GI enables chemical genetic screening of more than 13,000 compounds

Based on the encouraging results from our *in silico* evaluation, we designed a compressed mutant pool based on the COMPRESS-GI selection and used this as a basis for a large-scale chemical genetic screening effort (Fig 6a). The details of this effort and specific findings related to the screened compounds are described in our companion paper [35], but we briefly highlight the impact of the mutant selection here. Our chemical-genetic screening platform utilizes barcode sequencing technology, which leverages the fact that each mutant strain is barcoded with a unique 20bp identifier adjacent to a common priming site [34]. This design allows the selected set of mutants to be grown in a single pool for each compound interrogated [34]. Furthermore, depending on the number of unique strains being tracked, several different compound conditions can be multiplexed in a single sequencing lane, which can dramatically increase the throughput of the approach (Fig 6a). In our application, we constructed a mutant pool consisting of 157 genes selected by the COMPRESS-GI algorithm along with approximately 200 genes that were manually selected to complement the diagnostic set. This design enabled us to screen compounds at 200 uL and multiple 768 unique compound conditions per lane of Illumina sequencing. Using our diagnostic mutant pool, we achieved a similar per-strain read depth as genome-wide chemical-genetic screens at ∼15X higher throughput, saving significant resources (Fig 6b) in addition to smaller compound volumes.

**Fig 6.**
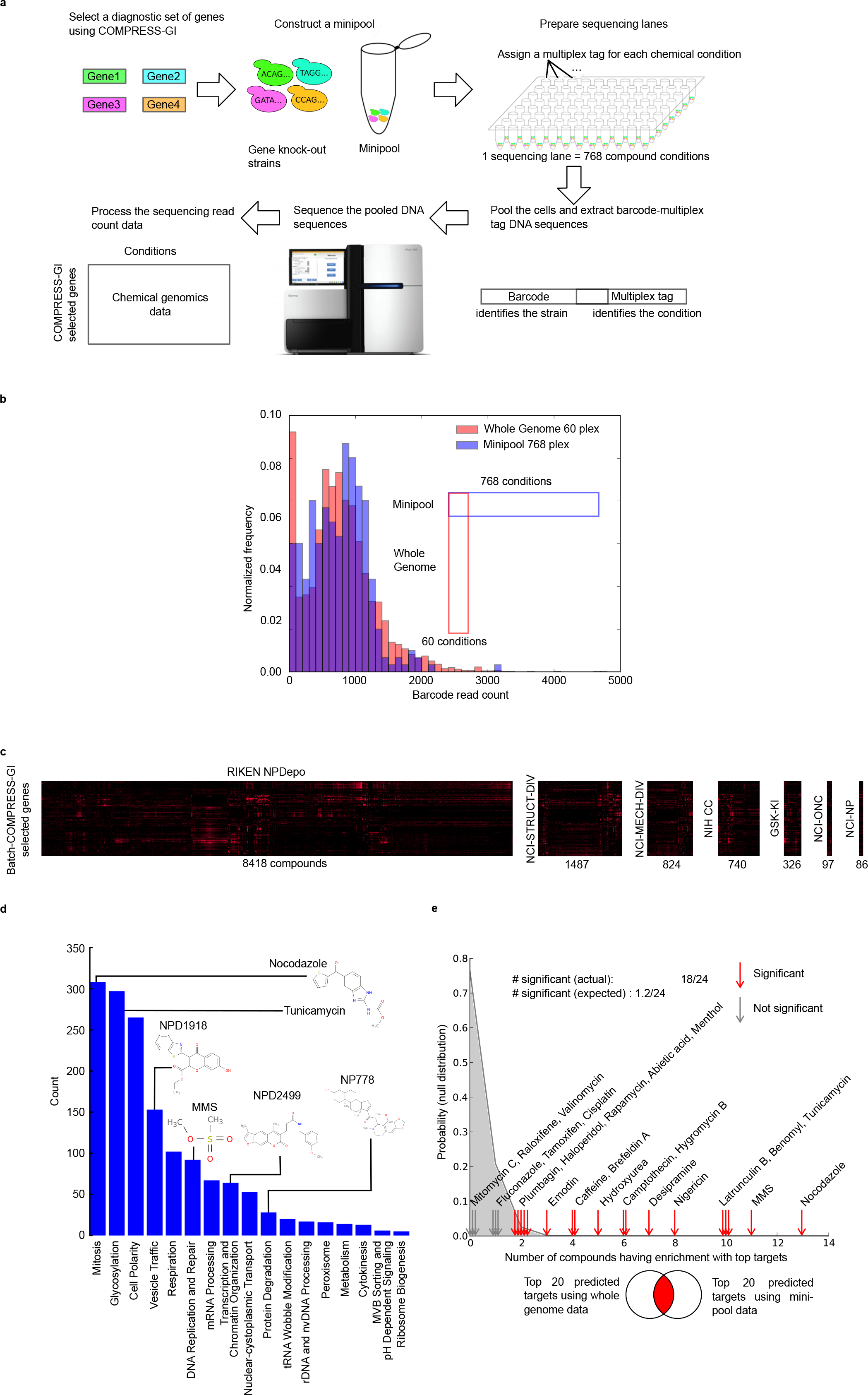
COMPRESS-GI enables large-scale chemical genetic screen and functional
interpretation of chemical genetic profiles. (a) The flowchart provides an overview of how COMPRESS-GI was used for the high-throughput chemical genetic screens in *S. cerevisiae*. COMPRESS-GI was applied to the *S. cerevisiae* genetic interaction data to identify a gene set that recapitulates a functionally predictive genetic interaction profile similarity network. The mutant strains corresponding to those selected genes were used to create a mutant pool, which was used to screen chemical genetic profiles using the barcode sequencing technology. (b) Using the COMPRESS-GI selected minipool, we were able to screen an order of magnitude more conditions (768 vs. 60) using same amount of sequencing resources, thereby reducing the sequencing cost for each condition. (c) Clustergram of the resulting chemical genetic data for 11,983 compounds across 7 major compound libraries screened using the COMPRESS-GI selected genes. (d) Distribution of the target process predictions for the high-confidence set of compounds in the screen across several biological processes. (e) The experiment shows the similarity of the top 20 targets predicted by Parsons *et al*. whole genome chemical genetic profiles as compared to predictions derived from the screens completed using the COMPRESS-GI designed diagnostic mutant set.

To validate the utility of our diagnostic mutant pool, we rescreened 24 compounds using our diagnostic mutant collection that were originally screened in the previous genome-wide study [31], which provided a direct opportunity to compare genome-wide profiles with compressed mutant profiles. For each of these compounds, we compared the top 20 most similar gene targets generated either from the diagnostic gene set or the genome-wide mutant collection. Indeed, we found the top predicted targets significantly overlapping between these two sets for 18 of the 24 compounds (Fig 6e), suggesting we are able to recapitulate predictions of mechanism of action using our compressed diagnostic pool.

In total, we screened more than 13,000 compounds across 5 different compound collections using this optimized screening approach enabled by COMPRESS-GI [35]. We were able to make high-confidence mode-of-action predictions for the targeted bioprocess through comparison to genetic interaction profiles for a total of more than 1500 compounds [35, 36], supporting the utility of compressed mutant profiles for functional characterization. The predicted modes of action spanned a diverse set of functional categories (Fig 6d), and we collected additional phenotypic data supporting our predictions for novel compounds in several different classes including tubulin inhibitors, cell cycle, and cell wall targeting compounds [35].

## Discussion

In this study, we demonstrated that genetic interaction screening efforts can be optimized. We developed and tested two related algorithms for optimal selection of mutants for genetic interaction screens, one addressing the scenario when a substantial base of screens already exists, and a second iterative algorithm that starts with no prior screen data. These algorithms both focused on the objective of measuring functional similarity based on interaction profile comparisons. Indeed, we were able to improve the efficiency of screens for this purpose in both scenarios. In general, our results suggest that if the goal of genetic interaction screens is to measure functional similarity, a relatively small collection of well-selected mutants (< ∼10% of the genome) is able to achieve much of the utility of larger profiles, which is an encouraging finding for genetic interaction screening efforts in other contexts. This compression is inherently tied to the modular structure of biological systems. Since most genes function as part of larger inter-dependent groups, the high-level structure of genetic networks can be explored without exhaustive screening of all mutants. This finding also has substantial utility in a variety of settings. For example, in our use of COMPRESS-GI to design an optimized chemical genetics platform, we achieved a ∼10-fold increase in throughput while requiring smaller compound volumes, which enabled systematic chemical genetic profiling of several large compound collections that, otherwise, would not have been possible.

We note that while we have focused our attention on optimizing genetic interaction screens for measuring functional similarity, there are other important objectives that motivate genetic interaction screening efforts. For example, in the context of human cells, detection of specific instances of synthetic lethality is of interest because each discovered individual synthetic lethal gene pair represents a potential target for personalized cancer therapy or modifier of a disease-associated mutation. If the objective is to most efficiently capture all possible cases of synthetic interactions, our analysis in yeast suggests this is a much more difficult setting to achieve substantial gains (S1a Fig). The compression achieved with our COMPRESS-GI algorithm for the profile similarity use case will not be possible to achieve in that setting, and more extensive screening efforts cannot be avoided. Another related objective is the exhaustive detection of specific between-pathway or within-pathway interaction structures that dominate genetic interaction networks [23]. Our analysis suggests that this is a similarly difficult objective that inherently requires substantially more screening than the profile similarity case (S1d Fig).

It is important to note that our algorithm relies on a supervised approach. We use knowledge of functional relationships between genes to guide the selection of mutants that enable the detection of functional similarity when interactions are measured with them. We have designed our approach to cover diverse functions and demonstrated that it works well even in cross-validation scenarios. However, we note that the Gene Ontology annotations we used as a basis for the functional relationship standard are more complete and of better quality in yeast than in many other settings. We anticipate there will be other settings where such a standard is lacking, which could limit the application of our approach. However, limitations in annotations may be compensated for by rich collections of unbiased genome-scale data. For instance, in the context of the human genome, we anticipate that current collections of human genomic data, coupled with methods to integrate them to form robust and even tissue-specific functional networks (e.g. [37]), could likely substitute for curation-based standards.

In general, we anticipate that strategies for efficient, yet systematic, screening of genetic interactions will continue to grow in importance. Genome-scale application of CRISPR-Cas9 genome editing approaches offer the potential to map large-scale genetic networks for a variety of different species, including in human cells. Larger, more complex genomes will necessitate rational data-driven strategies for exploring the enormous space of possible combinatorial perturbations. Even in yeast, where the first complete pairwise genetic interaction network now exists [3], there are new, important frontiers to be explored for which exhaustive mapping is simply not feasible. For example, moving from double mutants into the space of triple mutant combinations will require a more targeted approach, possibly guided by the concepts we describe. Recent studies also demonstrate the dynamic nature of genetic interactions across different environmental conditions [15, 38], which is an important dimension to be explored as well. The methods proposed here lay a foundation for how existing annotations, genomic data, and iteratively generated genetic interactions can be integrated to rationally explore genetic networks.

## Acknowledgements

This work was partially supported by the National Institutes of Health (R01HG005084, R01GM104975) and the National Science Foundation (DBI 0953881). CM and CB are fellows in the Canadian Institute for Advanced Research (CIFAR) Genetic Networks Program. Computing resources and data storage services were partially provided by the Minnesota Supercomputing Institute and the UMN Office of Information Technology, respectively. SWS was supported by an NSF Graduate Research Fellowship.

## Methods

### COMPRESS-GI

Given genetic interaction data (m query genes crossed against n array genes) and a Gene Ontology standard for the query genes (size m by m), the COMPRESS-GI method discovers an informative subset of array genes. The optimization objective for selecting the informative subset of genes is to maximize the match between the gene profile similarities based on the selected partial profiles and gene co-annotations in Gene Ontology. The matching is quantified using precision-recall statistics by treating gene profile similarities as predictions and co-annotations from Gene Ontology as the gold standard positive and unrelated genes in the Gene Ontology as gold standard negative. The informative set of genes is discovered by exhaustively searching for genes that when added to the selected set of genes will best improve the precision-recall statistics. For example, for discovering the first gene, we conduct an exhaustive search of all the array genes and the gene that gives the best precision-recall statistics is selected. For the second gene, we search for all the array genes except for the first selected gene, and select the gene that gives best precision-recall statistics along with the first gene. This process is continued until the precision-recall statistic saturates and the increase by adding any gene does not increase the precision-recall statistic significantly. In practice, a maximum of around 20 genes can typically be selected before the precision-recall statistic saturates.

The next set of genes to be discovered by the COMPRESS-GI is influenced by genes already selected by the algorithm. For example, different starting genes may result in convergence to a different final set. To ensure that the genes selected are robust to selection of the starting gene, we ran the COMPRESS-GI algorithm with different starting genes. For example, instead of starting with the best gene as the first gene, we started with the second best gene and allowed the first gene to occur in the COMPRESS-GI selections. We repeated this process with each of the 50 best genes ranked high in the precision-recall statistics based on the single gene profile as starting gene.

Further, to make sure that all the major functional categories are represented by the selected set of genes, we repeated the COMPRESS-GI algorithm for several different functional contexts. The functional context was created by limiting the Gene Ontology standard to only genes that are related to the function.

The different sets of genes obtained by running with different initial genes, and in different functional contexts, are combined, and the genes are sorted by their frequency of occurrence in these sets. The optimal number of the genes to be selected is determined based on the maximal precision at 25% recall averaged across different functional categories (see Fig 2e).

### Precision-recall statistic

The precision-recall statistic is a way to assess both precision as well as recall of predictions against a gold standard. In a typical machine learning setting, there is a positive and a negative class which are being predicted. If a positive prediction is found correct according to the positive gold standard, then the prediction is called True Positive otherwise it is called False Positive. Likewise, if a negative prediction is correct according to the negative gold standard it is True Negative otherwise it is False Negative. Precision is TP/(TP+FP) and Recall is TP/(TP+FN), where TP, FP, FN are number of True Positives, False positives, and False negatives predictions respectively.

In our case, precision-recall statistics are used to assess the match between the gene similarities based on partial profiles with co-annotations in Gene Ontology. To evaluate the predictions and compute precision we also need gold standard positive co-annotations and gold standard negative co-annotations. These gold standard co-annotation are generated from Gene Ontology using GRIFN[24] (see Creation of GO standard). The similarities are thresholded at different points (recall equal to integral powers of 2 and the last recall) where precision and recall statistics are calculated and the precision-recall curve is plotted. Since the denominator for recall is constant for all similarity thresholds (TP + FN = number of 1s in the GO standard matrix), we have ignored the denominator and used Recall = TP.

### Comparing Precision-recall curves

For the COMPRESS-GI approach, precision-recall curves are compared to find the best gene to select at each iteration. The precisions are compared at recall (TP) at powers of 2: 2, 4, 8, and so on. The precisions at earlier powers of 2 are compared first. If one of the PR curves has higher precision at that recall, that one is considered to be a better PR curve. In case of tie, precisions at higher recalls are considered. One problem with this approach is that after the PR curve has saturated, even weak profiles can become slightly better by chance. To safeguard against this situation, in addition to checking that the PR curve improves we also check that the increase is greater than the sum of standard error in the two precisions. Given precision p = TP/(TP+FP), where TP, FP are number of true positive and false positives, respectively, the standard error on p is calculated as 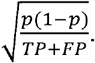

### Category specific precision-recall statistics

The COMPRESS-GI approach is run with several different functional contexts, that is, we want to select informative set of genes for the different functional category. To compute category specific precision-recall statistics and optimize on that objective, we modify the Gene Ontology standard to be specific to the functional category. The GO standard, M, is changed as follows:

(1) M_i,j_ is unchanged if genes i, j both belong to the functional
category,
(2) M_i,j_ = 0 if originally M_i,j_ == 1 and only one of the genes i, j belong to the functional category.
(3) M_i,j_ is unchanged if M_i,j_ = −1 originally

The GO standard for the genes within the functional category remain unchanged (1), but co-annotations of gene pairs outside the functional category are set to 0. Even though the focus of the optimization is to select genes informative for a particular functional context, the -1s in GO standard are never changed so that predicting unrelated genes as related is always penalized.

### iCOMPRESS-GI

Like the COMPRESS-GI, the iterative method is also based on a similar objective of optimizing the match between the similarities of the genes with Gene Ontology standard (***G***). The similarities of the genes based on the partial profiles can be written as ***XW*(*XW*)^*T*^ = *XWW^T^X^T^ = XWX^T^*** where ***W*** is the diagonal matrix with ***W_ii_* = 1** if array gene i is selected. However, unlike the COMPRESS-GI where precision-recall statistics are used to assess the match between ***XWX^T^*** and ***G***, we optimize on the sum of element wise multiplication of ***XWX^T^*** and ***G.*** This objective can be written as

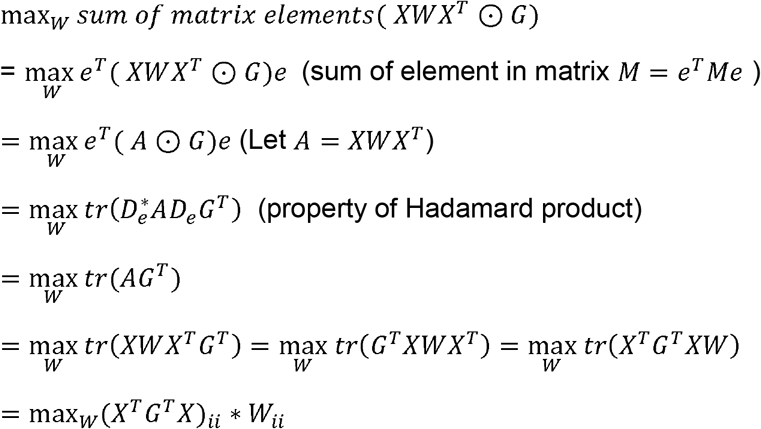

where ʘ is the element wise multiplication and more formally known as Hadamard product, and *tr* refers to the trace of a matrix which is sum of diagonal elements.

This reduces the problem to the simplest type of 0-1 knapsack problem which can be solved by a greedy algorithm. To solve this problem, we rank the genes by (*X^T^G^T^X)_ii_*, and pick the top n genes.

#### Algorithm Complexity

The complexity of the iCOMPRESS-GI algorithm mainly lies in the matrix multiplication *X^T^G^t^X*. So if *X* is the genetic interaction matrix composed of *m* queries and *n* arrays, the complexity for *X^T^G^T^X* matrix multiplication is O(*nm*^2^+ *mn*^2^) = O((*mn*)(*m* + *n*)). The complexity of the knapsack problem is O(*n*), so the overall complexity of the algorithm is O((*mn*)(*m* + *n*)). This complexity makes the algorithm perfectly reasonable to run on genetic interaction datasets that are several folds larger than the current largest genetic interaction datasets [3]. Further, the algorithm can be used even for organisms with a much larger number of genes (*m,n* ∼ 60,000). This complexity allows the algorithm to be run very quickly for iterative approaches, which has been specifically demonstrated in the results.

### Interaction profile similarity measure

We have used the dot product similarity measure because we showed earlier that it is among the best performers on genetic interaction data for predicting gene associations [39]. More importantly, similarity measures that use normalization such as Pearson correlation, Cosine correlation and Spearman correlation are unstable on smaller profiles, so they could not be used for this study where we are building the informative set of genes. Further, dot product has been shown to be more robust to noise and batch effects, which are typical in genetic interaction data [39].

### Block discovery

To discover block structure in the genetic interaction data, we have used a block discovery method published earlier by our lab, XMOD [23]. We streamlined the XMOD implementation and provide a python interface to XMOD so that it can be run from the command line. For all block discovery applications (Fig 1d), we use a minimum support threshold of 6 and item-set size of 3, which means all the discovered blocks are greater than or equal to size of 6 (on the query side) by 3 (on the array side). The blocks discovered are compared with blocks obtained by running XMOD on degree distribution preserved randomized genetic interaction network (see Bellay *et al.[23])*, and we use a p-value of 0.001 to filter out insignificant blocks. The discovered set of blocks may be overlapping, so we remove blocks which share an overlap of 10% or more with a larger block.

### Gene Ontology

The *S. cerevisiae* and *S. pombe* Gene Ontology (GO) standard were created using the approach described in [24], which generates a co-annotation matrix based on the *S. cerevisiae* Gene Ontology [40] and annotations. The S. cerevisiae datasets were downloaded on January 22, 2012, and the *S. pombe* datasets were downloaded on July 4, 2012. The final standard includes annotations for all pairs of genes with some denoted as positives (functionally related), some as negatives (not functionally related), and some as zero (neither). The S. *S. cerevisiae* standard contains 5513 genes while the *S. pombe* dataset contains 4598 genes. The GO standards for both species are available for download on the Supplementary website (http://csbio.cs.umn.edu/compress-gi).

### Genetic Interactions

We used the *S. cerevisiae* and *S. pombe* genetic interaction datasets from Costanzo et al. [4] and and Ryan et al [27] studies, respectively. These datasets contain multiple alleles or multiple conditions for some strains which can confound some of the analyses in our study. In such cases, only the strain/condition profile that had the highest negative genetic interaction degree (genetic interaction ε < −0.08, pval < 0.05) was retained. The resulting data sets contain 1672 query (rows) by 3885 array (columns) genes for *S. cerevisiae* dataset and 879 query (rows) genes by 1955 array (columns) genes for *S. pombe* dataset.

For parts of our evaluation of COMPRESS-GI and iCOMPRESS-GI, square genetic interaction matrices were required for each species where genes on the row side are also on the column side. These square genetic interaction matrices were derived from the rectangular matrices for both the species by retaining only the common genes for both row and column sides. The *S. cerevisiae* square matrix contains 1141 rows and columns and the *S. pombe* square matrix contains 408 rows and columns.

### Source Code

The source code for the methods are available at: http://csbio.cs.umn.edu/compress-gi.

---

**Supplementary Fig 1. Evaluation of baseline strategies for common genetic interaction screening objectives.** (a) Genetic interaction coverage: The figure shows how selection strategies of random genes (gray) and genes prioritized based on single mutant fitness defects affect the percentage of the interactions covered. The random selection was conducted 21 times, and the light gray region is the inter-quartile range while the dark gray line is the median across random selections. Similar analyses are repeated for the other genetic interaction use cases. (b) Profile similarity network: The functional information in the profile similarity network was measured using precision-recall analysis (see Online Methods). Precision is sampled at a recall of 2048 true positives (∼2% recall). (c) Genetic interaction degree estimation: The genetic interaction degree is estimated with partial profiles, and Pearson correlation is used to compare with actual degrees obtained from the complete dataset. (d) The number of bipartite or the block structures in the genetic interaction network: The number of significant blocks was discovered by running XMOD [23] with different sets of array genes.

**Supplementary Fig 2. Genetic interaction information is concentrated in the top few principal components, suggesting potential for compression**. Standard PCA analysis was conducted to estimate the number of unique principal components explaining the majority of observed variation in the genetic interaction data. The figure plots the cumulative fraction of variance explained (y-axis) captured by the top most informative components (x-axis). Consistent with our findings, this analysis suggests the possibility of substantial compression given the disproportionate among of variance explained by the early principal components.

